# Non-bursting non-rhythmic neurons of the ventral pallidum form cell assemblies and respond to reward and punishment during Pavlovian conditioning

**DOI:** 10.1101/2020.04.21.053843

**Authors:** Panna Hegedüs, Julia Heckenast, Balázs Hangya

## Abstract

The ventral pallidum (VP) is a major hub interfacing striatopallidal and limbic circuits, conveying information about salience and valence crucial to adjusting behavior. However, how neuron populations of the VP with different firing properties represent these variables is not fully understood. Therefore, we trained mice on auditory Pavlovian conditioning and recorded the activity of VP neurons while mice were performing the task. Many VP neurons responded to punishment (51%) and reward (44%), either by firing rate increase or decrease. Additionally, 20% of cells responded to outcome-predicting auditory stimuli, showing larger responses to reward-predicting cues compared to those that signaled likely punishment. We found that a large subset of VP neurons showed burst firing based on their auto-correlograms, while a small population exhibited fast rhythmic discharge in the beta/gamma frequency range. Some bursting neurons exhibited distinct response properties of their bursts and single spikes, suggesting a multiplexed coding scheme in the VP. However, non-bursting, non-rhythmic neurons were the most sensitive to reward and punishment. Finally, we demonstrate the presence of synchronously firing neuron assemblies in the VP. Neurons participating in such assemblies were particularly responsive to reinforcing stimuli. This suggests that a synchronous, non-bursting, non-rhythmic neuron population of the VP is responsible for the lion’s share of ventral pallidal salience representation, likely important for reinforcement learning.

**Significance statement:** The ventral pallidum (VP) is a subcortical brain area that participates in regulating motion and emotion by processing information related to appetitive and aversive stimuli. However, how these stimuli are represented by VP neural circuits is not well understood. Therefore, we investigated how VP neuron populations defined by their firing properties respond to reward and punishment during Pavlovian conditioning. We found that a distinct, non-bursting-non-rhythmic group of neurons was responsible for most responses to reward and punishment in the VP. Neurons of this group formed co-active cell assemblies and multiplexed different types of information via different firing patterns, revealing flexible and plastic neuronal representation strategies in the VP during associative learning.

## Introduction

The VP serves as an interface between the limbic system and other structures, integrating cortical, amygdala, basal ganglia and neuromodulatory input. On the effector side, its projections to thalamus, cortex, basal ganglia and other subcortical structures including hypothalamus, ventral tegmental area (VTA) and lateral habenula (LHb) influence motivation, reinforcement learning and attention (Zahm et al., 1996; Maurice et al., 1997; Ito and Doya, 2009; Smith et al., 2009; Root et al., 2015; Richard et al., 2016a; Faget et al., 2018; Stephenson-Jones et al., 2020). Specifically, crucial aspects of ventral pallidal activity in associative learning have been revealed recently, showing VP neurons encoding incentive salience and valence as well as mounting behavioral response to environmental changes (Tindell et al., 2004, 2006; Avila and Lin, 2014; Richard et al., 2016a; Stephenson-Jones et al., 2020).

Recent research has focused on disentangling the role of VP GABAergic and glutamatergic projection neurons in signaling aversive and appetitive stimuli. A consensus is emerging, postulating that positive and negative value signals are transmitted to VTA and LHb via inhibitory and excitatory projection neurons of the ventral pallidum, respectively (Knowland et al., 2017; Faget et al., 2018; Wulff et al., 2019; Stephenson-Jones et al., 2020). Moreover, these projections may differentially drive reward seeking and threat avoidance, bearing strong relevance to drug-seeking, relapse and depression (Knowland et al., 2017; Prasad et al., 2020; Stephenson-Jones et al., 2020).

At the same time, the VP lies at the border of the basal ganglia and basal forebrain (BF) circuitry, housing cholinergic neurons usually interpreted as part of the BF (Záborszky and Cullinan, 1992; Zaborszky et al., 2012; Root et al., 2015). Indeed, VP has been shown to simultaneously convey movement and reward-related information (van den Bos and Cools, 1991; Tachibana and Hikosaka, 2012; Avila and Lin, 2014; Richard et al., 2018; Stephenson-Jones et al., 2020), both prominently represented by BF neurons as well (Lin and Nicolelis, 2008; Fuhrmann et al., 2015; Hangya et al., 2015). Thus, the VP has to multiplex different outcome-related information from a set of sources that serve overlapping but different functions including approach and avoidance, motivation, movement vigor, learning and memory.

How are different types of information encoded by the VP? Whether they are routed through different lines of this intricate switch board, labelled by markers such as parvalbumin, the vesicular glutamate transporter VGLUT2 or the inhibitory marker GAD2 has been explored recently (Knowland et al., 2017; Faget et al., 2018; Prasad et al., 2020; Stephenson-Jones et al., 2020). However, another exciting possibility is that integrating and multiplexing is also represented by different coding schemes including elements of rate and temporal code, such as characteristic firing patterns like bursts or single spike firing, rhythmic discharges, network level synchrony and asynchronous activity. Accordingly, Avila and Lin suggest that electrophysiological characterization that goes beyond the broad categories of inhibitory and excitatory cell types will enable a better understanding of how VP performs its functions (Avila and Lin, 2014).

To address the above question, we recorded VP neurons while mice performed a probabilistic Pavlovian conditioning task. By using auto- and cross-correlation techniques, we uncovered the presence of separate fast rhythmic, bursting and non-bursting-non-rhythmic neurons, similar to previous electrophysiological categorization of VP neurons (Pang et al., 1998). We found that reinforcement-related signals were most frequent in the non-bursting, non-rhythmic population. The analysis of synchronous discharges revealed the presence of co-firing neuron assemblies. Cells that participated in these synchronously firing assemblies showed increased responsiveness to reinforcers. Thus, VP neurons with distinct discharge types both at the individual and network level, likely corresponding to different coding strategies, show differences in their representation of behaviorally salient stimuli. Even within single neurons, burst and single spike firing could strongly dissociate, suggesting the presence of a distinct burst code in the VP (Kepecs et al., 2002; Kepecs and Lisman, 2003).

## Methods

### Animals

Adult male mice (n = 3 ChAT-IRES-Cre, B6129F1 and n = 1 PV-IRES-Cre, FVB/AntFx) were used according to the regulations of the European Community’s Council Directive of November 24, 1986 (86/609/EEC). Experimental procedures were reviewed and approved by the Animal Welfare Committee of the Institute of Experimental Medicine, Budapest and by the Committee for Scientific Ethics of Animal Research of the National Food Chain Safety Office of Hungary.

### Surgery

Mice were anesthetized with an intraperitoneal injection of ketamine-xylazine (0.166 and 0.006 mg/kg, respectively) after a brief induction with isoflurane. After shaving and disinfecting the scalp (Betadine), the skin was infiltrated with Lidocaine and the eyes were protected with eye ointment. Mice were placed in a stereotaxic frame and the skull was levelled along both the lateral and the antero-posterior axes. The skin, connective tissues and periosteum were removed from the skull and a cranial window was drilled above the anterior ventral pallidum (antero-posterior 0.75 mm, lateral 0.6 mm). Two additional holes were drilled above the parietal cortex for ground and reference. After virus injection to the VP (AAV 2/5. EF1a.Dio.hChR2(H134R)-eYFP.WPRE.hGH), a custom-built microdrive (Kvitsiani et al., 2013; Hangya et al.,2015) was implanted into the VP using a cannula holder on the stereotactic arm. The microdrive and a titanium headbar were secured to the skull with dental cement (LangDental acrylic powder and liquid resin, C&B Metabond quick adhesive cement). The analgesic buprenorphine (Bupaq) was administered and mice were allowed a 1-week recovery period and handled for an additional week before training and recording.

### Pavlovian cued outcome task protocol

Mice were trained on an auditory Pavlovian conditioning task in a head-fixed behavioural setup described in detail previously (Solari et al., 2018). On the first day of training, thirsty mice were head-fixed and given free access to water reward whenever they licked a waterspout. The next day, a pure tone cue was introduced that predicted likely reward. After each cue presentation, water reward was delivered with 0.8 probability with a 400-600 ms delay, while the rest of the outcomes were omissions. Next, a second pure tone cue of well-separated pitch was introduced that predicted reward with low probability (0.25). Air puff punishment was introduced in the following session with the final outcome contingencies (likely reward trials, 80% reward, 10% punishment, 10% omission; likely punishment trials, 25% reward, 65% punishment, 10% omission). The trials with different trial types (likely reward and likely punishment) and outcomes (water reward, air puff punishment and omission) were presented in a pseudorandomized order. Mice learned the task in approximately one week and from the second week, consistently demonstrated reward anticipation by differential licking rate in response to the cues (Fig.1E-H).

**Figure 1.**
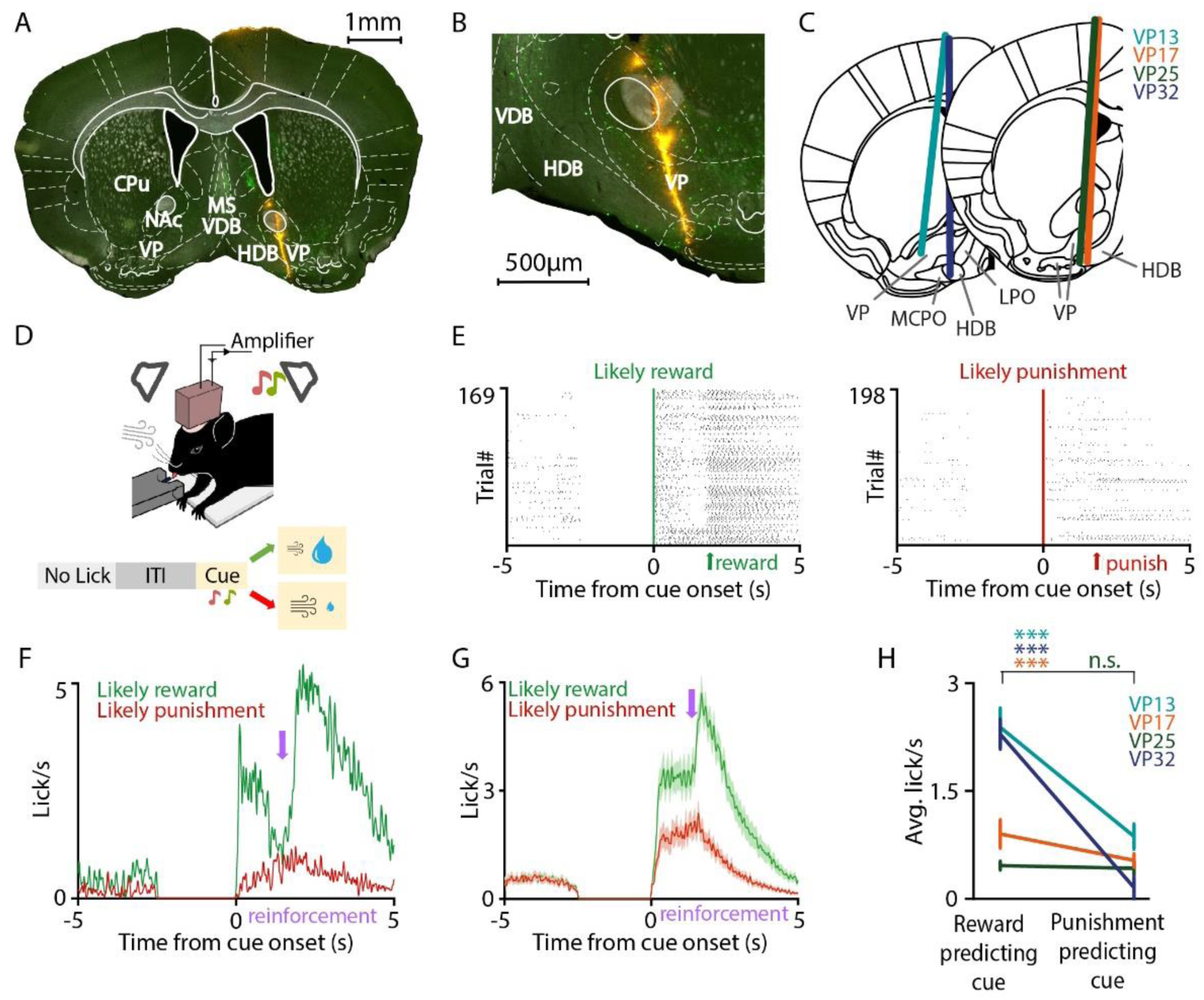
Targeting VP in mice performing auditory Pavlovian conditioning. ***A***, Coronal section from a ChAT-Cre mouse showing the tetrode tracks (DiI, red; ChAT+, green) through the VP. ***B***, Magnified view showing viral expression of ChR2-eYFP in the target area. ***C***, Reconstructed location of the electrode tracks. ***D***, Top, schematic of the auditory Pavlovian task setup. Bottom, trial structure with possible outcomes. After the mouse stopped collecting the previous reward (‘no lick’), a variable inter-trial interval started, signaled by turning an LED off, in which no licking was allowed. Then two cue tones of well-separated pitch predicted likely reward or likely punishment. ***E***, Raster plots of licking activity in an example session of one mouse. The cue predicting likely reward (left, green) elicited stronger anticipatory licking than the cue that signaled likely punishment (right, red). ***F***, Peri-event time histograms (PETH) of licking activity of the same session. ***G***, Average licking activity (PETH) of N = 4 mice shows stronger anticipatory licking after the cue that predicted likely reward (green). Purple arrow indicates average reinforcement delivery time. ***H***, The reward-predicting cue elicited significantly more licks in 3/4 mice. CPu, caudate putamen; HDB, horizontal nucleus of the diagonal band of Broca; MS, medial septum; NAc, nucleus accumbens; VDB, vertical nucleus of the diagonal band of Broca.

### Recording

Extracellular recordings were performed with custom made microdrives consisting of 8 movable tetrode electrodes and an optical fiber. The microdrive was connected (Omnetics) to a 32-channel RHD headstage (Intan). Data were digitized at 32 kHz and transferred from the headstage to a data acquisition board (Open Ephys) via a Serial Peripheral Interface cable (Intan).

### Data analysis

Data analysis was carried out using custom written Matlab code (Mathworks). Action potentials were sorted into putative single neurons manually by using MClust (A.D Redish). Only neurons with good cluster quality (isolation distance > 20 and L-ratio < 0.15) were included in the final dataset for further analysis (Schmitzer-Torbert et al., 2005; Hangya et al., 2015).

After spike sorting, the activity of individual neurons was aligned to different task events (cue presentation, reward and punishment delivery). Statistics were carried out on each neuronal unit; a baseline activity was defined by taking a 1 s window before the cue, then firing rate in the baseline window was compared to firing rate in the test window (0-0.5 s after the event) by Mann-Whitney U-test (p < 0.001). Neurons were sorted into different groups based on their statistically significant responses to these events (e.g. activated by cue, inhibited by reward etc.).

Auto-correlograms (ACG) were calculated at 0.5 ms resolution. *Burst index* (BI) was calculated by the normalized difference between maximum ACG for lags 0-10 ms and mean ACG for lags 180-200 ms, where the normalizing factor was the greater of the two numbers, yielding an index between −1 and 1 (Royer et al., 2012). A neuron with a BI > 0.2 was considered to be bursting based on empirical observation reported previously (Laszlovszky et al., 2020) and confirmed by the presence of ‘burst shoulders’ on average ACG in the ‘bursting group’ and the complete lack of ‘burst shoulder’ on the average ACG in the ‘non-bursting’ group. To examine burst coding in the VP, analysis of neuronal responses to reinforcement-predicting cues and reinforcers were also carried out when only bursts or single spikes were considered for a neuron. A burst was detected whenever an inter-spike interval (ISI) was < 10 ms and subsequent spikes were considered as part of the burst as long as the ISI remained < 15 ms.

Characterization of rhythmic firing in the beta-gamma range was performed based on autocorrelograms. ACG peaks were detected either in the beta (16-30Hz) or gamma (30-100 Hz) frequency range. Then, the average value of a small window (±20 ms) around the peak was compared to a value calculated from a baseline period with the same algorithm. Neurons were considered rhythmically firing when this ratio was > 0.4 for the beta and > 0.25 for the gamma band. These cutoff values were determined empirically and confirmed by observing all ACGs after sorting into rhythmicity groups.

Cross-correlograms (CCG) were calculated at 1 ms resolution. CCGs were calculated and plotted for all simultaneously recorded pairs of neurons. Synchronously activated pairs were sorted based on a significant peak exceeding the upper 95% confidence interval by at least 10 counts of co-occurrences in the CCG around zero lag. A 1-2 ms wide asymmetric peak between 1-4 ms time lags was considered a putative monosynaptic excitatory connection based on previous reports (Bartho et al., 2004; Fujisawa et al., 2008; Hangya et al., 2010). Zero-lag synchrony was not tested for pairs of neurons recorded by the same tetrodes due to potential cluster contaminations during spike sorting.

For plotting average ACG and CCG, data were Z-score normalized with their surrogate mean and standard deviation. The surrogates were generated using the shift predictor method that introduces randomized delays between the correlated signals to generate a null distribution of no correlated activity (Fujisawa et al., 2008).

### Experimental design and statistical analyses

This study includes the analysis of 573 neurons recorded from 4 mice. These sample sizes were determined according to the standards of the field and exceed the minimal requirements of most statistical tests. However, this strategy is necessary as subsequent statistics after sorting neurons into groups have group sample sizes that are not possible to plan before conducting the experiments.

Statistical comparison of central tendencies was performed using non-parametric tests (Mann-Whitney U-test for unpaired data and Wilcoxon signed rank test for paired data) as normal distribution of the underlying data could not be determined unequivocally. Distributions over categorical variables were compared by chi square test for homogeneity. The exact p-values were reported for group comparisons.

Significant firing rate changes were evaluated at p<0.001 (Mann-Whitney U-test) to keep false positive rate low. Significant activation in cross-correlograms was determined by 95% confidence intervals generated by the shift predictor method (Fujisawa et al., 2008; Kvitsiani et al., 2013). We introduced a lower bound on effect size and required >=10 counts above this limit to disregard very small effects, which also ensures the robustness of the bootstrap process of surrogate generation. Average auto- and cross-correlations were calculated using Z-score normalization based on a surrogate null hypothesis distribution as described above, to allow equal weighting of individual neurons in the average.

### Code accessibility

MATLAB code developed to analyse the data presented in this study is available at www.github.com/hangyabalazs/VP_data_analysis.

### Histology

After the *in vivo* experiments, animals were anesthetized with an intraperitoneal injection of ketamine-xylazine (0.166 and 0.006 mg/kg, respectively) and transcardially perfused with saline for 2 minutes and 4% para-formaldehyde for 20 minutes. The brain was removed from the skull and 50 µm sections were cut using a vibrotome (Leica). The sections were mounted on microscopy slides in Aquamount mounting resin. Fluorescent micrographs of the sections were taken using a Nikon C2 confocal microscope.

Track reconstruction was carried out by registering experiment logs with the fluorescent images (Hangya et al., 2015). The images were aligned to the Mouse Brain Atlas (Paxinos et al., 2001) to determine the recording locations.

## Results

### Ventral pallidal neurons are modulated by salience, valence and expectation during Pavlovian conditioning

To test how different ventral pallidal neurons represent behaviorally relevant events during classical associative learning, we trained mice (N = 4) on a probabilistic auditory Pavlovian conditioning task and monitored the activity of VP neurons (n = 573) (Fig. 1A-C). Mice were water restricted and head-fixed for training, listening to two pure tones of different pitch, where one tone predicted likely water reward (80% reward, 10% punishment, 10% omission) and the other tone predicted likely punishment (25% reward, 65% punishment, 10% omission; Fig. 1D). Mice learned to discriminate the cue tones, indicated by differential licking activity after cue onset in anticipation of reward (Fig. 1E-H).

During the task, 60% of VP neurons showed phasic, short latency activation or inhibition after at least one type of behaviorally salient stimuli of the task, that is, reward, punishment and/or the reinforcement predicting cues (Fig. 2A-F). Around 20% (n = 117/573) of the neurons were modulated by the cues, 44% (n = 250/573) by reward and 51% (n = 263/515, neurons recorded early in training could not be tested; Mann-Whitney U-test, p < 0.001) by punishment (Fig. 2G-L). The majority of significantly responsive neurons showed activation, with a smaller fraction of inhibited cells (activated by cue, 63/117, 54%; reward, 180/250, 72%; punishment, 194/263, 74%). Interestingly, there was a difference in the latency of peak activation or inhibition across responses to cue, reward or punishment. VP neurons responded the fastest to punishment, intermediate to reward and slowest to outcome predictive sensory cues (Fig. 2M-N).

**Figure 2.**
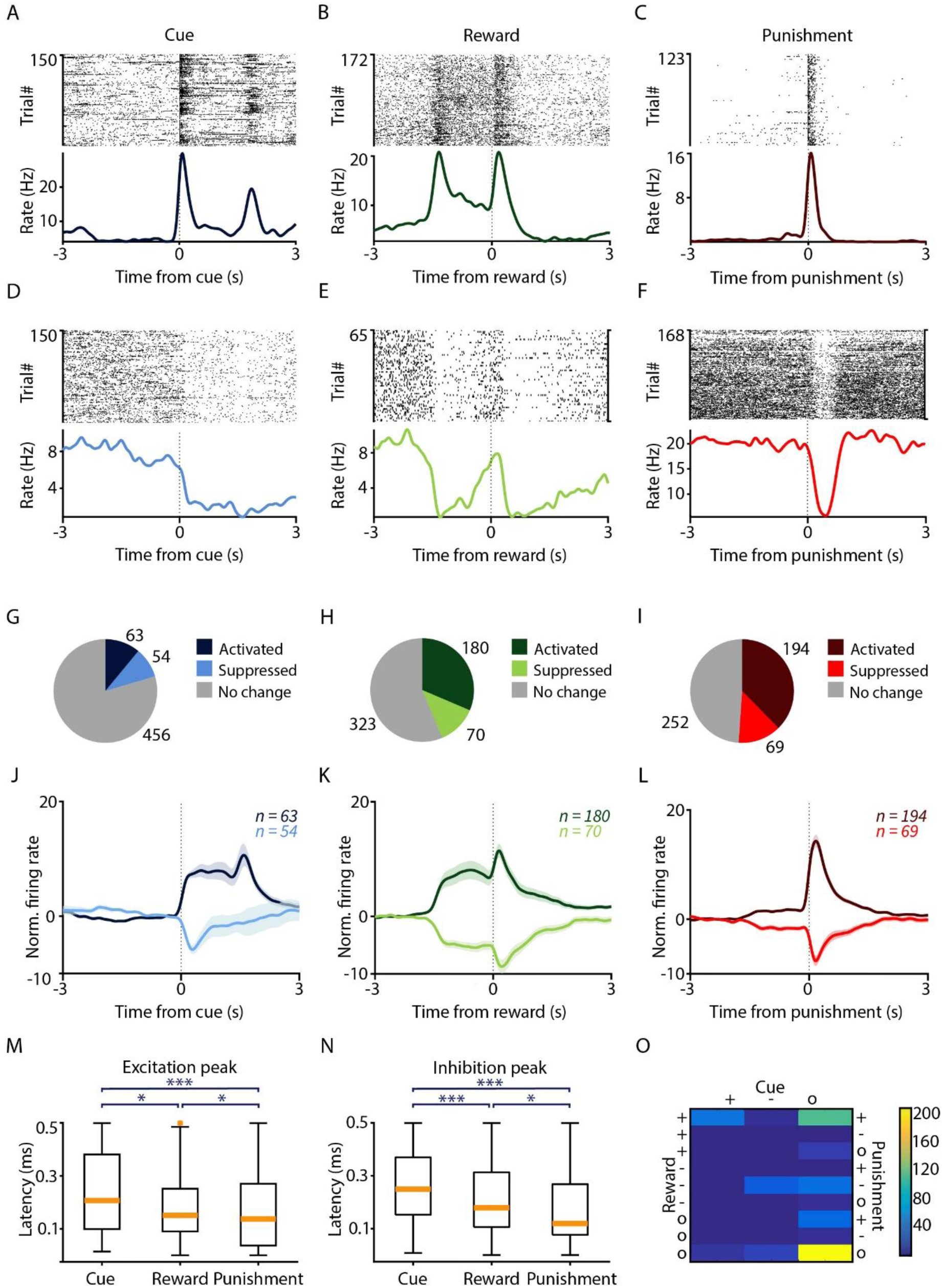
Ventral pallidal neurons are modulated by reward, punishment and outcome-predictive cues during Pavlovian conditioning. ***A-F***, Example VP neurons activated by cue stimuli (***A***), reward (***B***) or punishment (***C***), or inhibited by cue (***D***), reward (***E***) or punishment (***F***). Top, spike raster; bottom, PETH aligned to the respective behavioral events. ***G-I***, Pie charts showing the number of VP neurons activated and inhibited by cue (***G***), reward (***H***) and punishment (***I***). ***J-L***, Average z-scored PETH of VP neurons modulated by cue (***J***), reward (***K***) and punishment (***L***). Shading indicates SEM. ***M***, Excitatory response latencies to predictive cues, reward and punishment. ***N***, Inhibitory response latencies to predictive cues, reward and punishment. ***O***, Number of neurons showing different combinations of all possible responses to cues and reinforcers.

We found that neural responses to reinforcement of opposite valence were often correlated. For instance, in 158/515 neurons, the same neuron responded with increased firing rate to both the positive and negative reinforcer. Similarly, 59/515 neurons showed firing rate decrease after both reward and punishment (Fig. 2O). In contrast, only n = 2/515 neurons showed opposite responses to reward and punishment. Additionally, a large fraction of neurons showed correlated responses to reward predictive cues and primary reward (n = 44/515 activation and n = 24/515 inhibition to both, respectively). We have found a strong co-occurrence of activation to all behaviorally salient events including cue, reward and punishment (n = 41/515).

In sum, VP neurons showed an array of responses to behaviorally salient events. The fastest and most prevalent response pattern was a rapid activation after punishment. Direction of firing rate modulation was correlated across behaviorally relevant events, suggesting salience coding by individual VP neurons.

The probabilistic nature of the Pavlovian task meant that different cues were followed by reward, punishment or omission with different (but fixed) contingencies. Mice learned these contingencies (Fig.1), which required integration of positive and negative outcomes over many trials. Based on the predictive auditory cues we played before the reinforcement, reward and punishment could be either expected or surprising according to task contingencies. This allowed us to compare VP neuronal responses to expected vs. surprising outcomes.

We found that VP neurons strongly differentiated the distinct predictive cues, showing larger responses to cues that predicted likely reward (Fig.3A-B). Additionally, VP neurons showed slightly but significantly stronger responses to expected vs. surprising reward (Fig.3C-D). In contrast, VP neurons showed a tendency of responding more to surprising than expected punishment (Fig.3E-F). Taken together, VP responses were larger in those trials when the cue signaled likely reward, after which the delivery of reward was expected, while encountering an air puff punishment was surprising. This is consistent with the notion that VP responses may be scaled by incentive salience, but at odds with a role in traditional reward prediction error coding.

**Figure 3.**
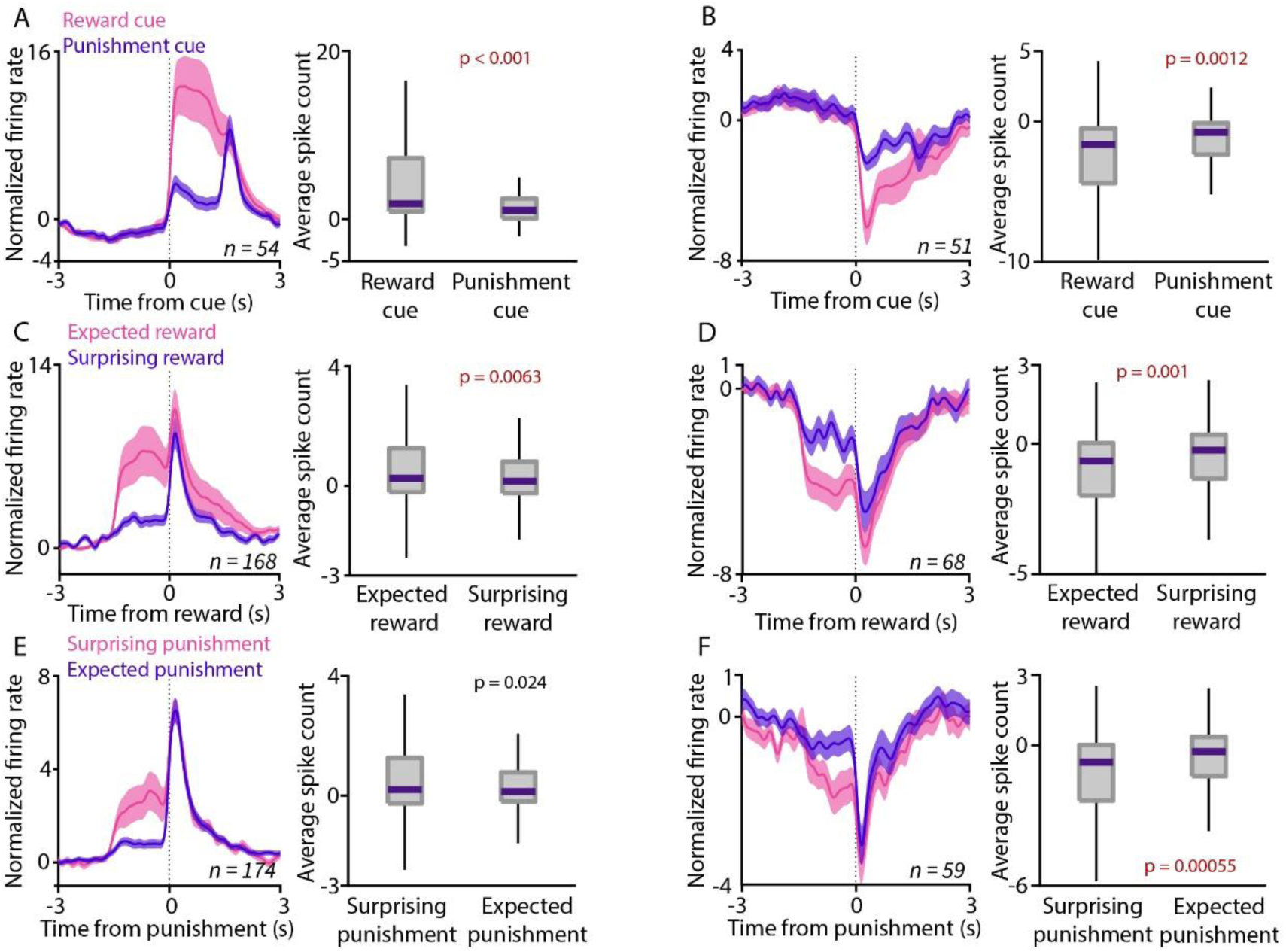
Ventral pallidal neurons are modulated by expectation. ***A***, Left, average PETH of VP neuronal activation after cues predicting likely reward (pink) or likely punishment (purple). Right, box-whisker plot comparing average spike count with respect to baseline after the two cues in VP neurons activated by cue presentations. ***B***, Average PETH and spike count relative to baseline for VP neurons inhibited after cue presentations. ***C-D***, Same as in ***A-B***, but showing responses to reward presentations. Pink, expected reward; purple, surprising reward. ***E-F***, Same as in ***A-B***, but showing responses to punishment presentations. Pink, surprising punishment; purple, expected punishment.

### Non-bursting, non-rhythmic VP neurons are more responsive to behaviorally salient events

Burst coding of salient events has emerged as a general scheme for subcortical representations: burst responses to behavioral reinforcement and reward predictive stimuli has been demonstrated for the VTA (Schultz et al., 1997), striatum, basal forebrain (Lin and Nicolelis, 2008; Hangya et al., 2015) and lateral habenula (Yang et al., 2018). To test whether this principle generalizes to the ventral pallidum, we categorized VP neurons as bursting and non-bursting based on short-latency peaks in their spike auto-correlations, indicating preferential firing with short inter-spike intervals characteristic of bursts (Royer et al., 2012; Laszlovszky et al., 2020) (Fig.4A; Methods).

**Figure 4.**
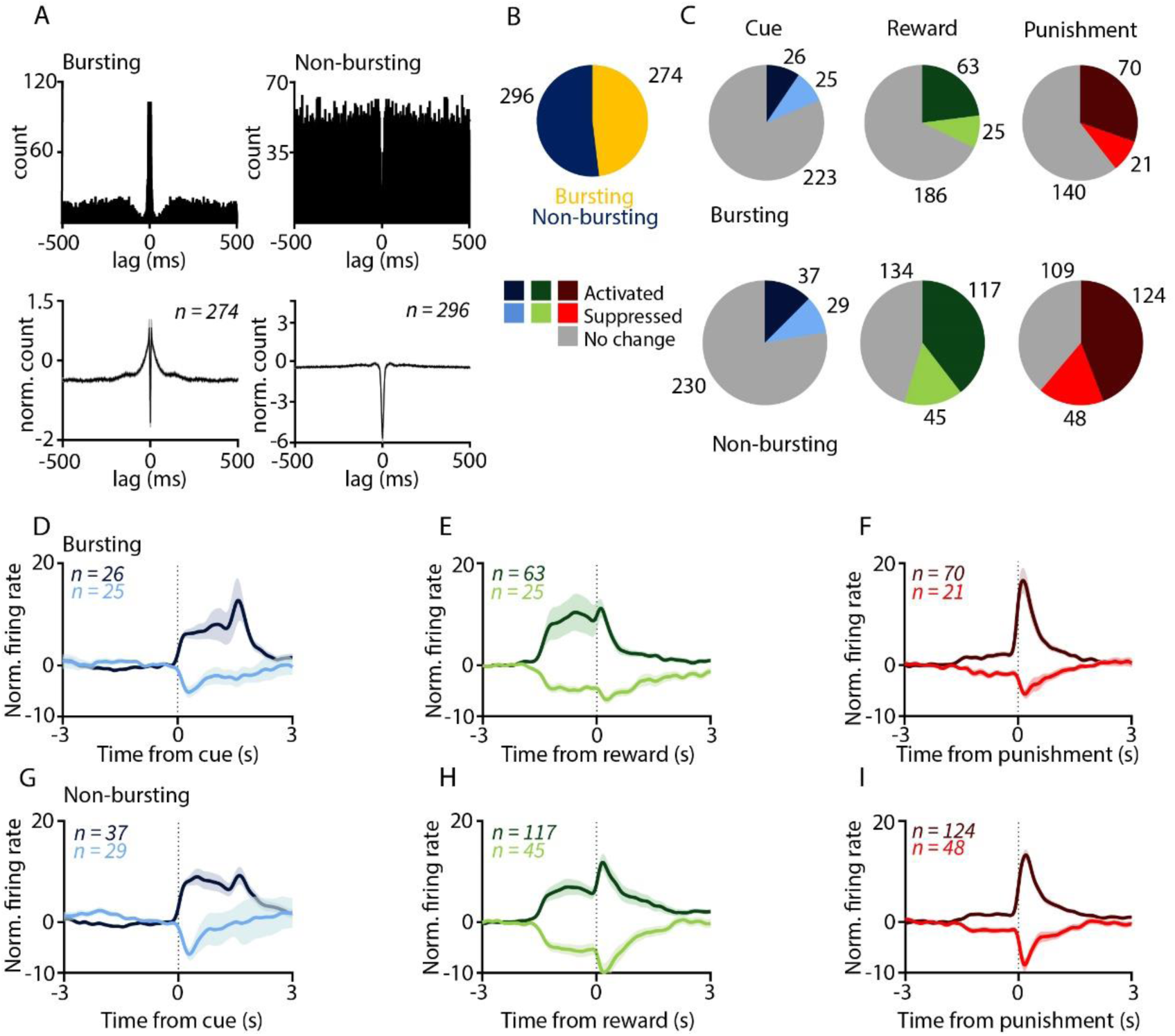
Non-bursting ventral pallidal neurons are more responsive to reinforcers. ***A***, Top, Example auto-correlograms of a bursting and a non-bursting VP neuron. Bottom, average. ***B***, Pie chart showing the proportion of bursting and non-bursting neurons. ***C***, Pie charts showing the number of bursting and non-bursting VP neurons activated and inhibited by cue, reward and punishment. ***D-I***, Average, z-scored PETHs of bursting ***(D-F)*** and non-bursting ***(G-I)*** VP neurons aligned to cue ***(D,G)*** reward ***(E,H)*** and punishment ***(F,I).***

We found that about half of VP neurons (n = 274 / 570, 48%; 3 neurons firing <100 spikes excluded from this analysis) fired bursts, while the remaining neurons were categorized as non-bursting (n = 296/ 570, 52%, Fig 4B.). Next, we tested whether bursting neurons were more responsive to reward, punishment and reward-predictive cues in Pavlovian conditioning. Surprisingly, we found that a larger fraction of non-bursting VP neurons showed significant responses to reinforcement (chi square test, p = 5.45 × 10^−8^ and p = 2.56 × 10^−9^ for reward and punishment, respectively, Fig 4C-I.). This was consistent both for reward and punishment, with a higher number of non-bursting neurons showing either firing rate increase or decrease. We did not find any difference regarding the fraction of cue-responsive neurons (chi square test, p = 0.2766).

Based on rhythmic modulation of their auto-correlation functions, we detected a small subset of rhythmically firing VP neurons (n = 38/573, 7%; Fig. 5A-B). We estimated the frequency at which these neurons were oscillating by their auto-correlation peak location and found that they fell in the beta/gamma range (6 beta-rhythmic and 32 gamma-rhythmic neurons were detected; Methods). These neurons have previously been identified as somatostatin-expressing GABAergic neurons (Espinosa et al., 2019). We found that most of these rhythmically discharging neurons showed weak or no responses to behaviorally salient events including cue tones, reward and punishment (Fig. 5C-I). These results suggest that mostly non-bursting, non-rhythmic neurons are recruited during reinforcement learning in the VP.

**Figure 5.**
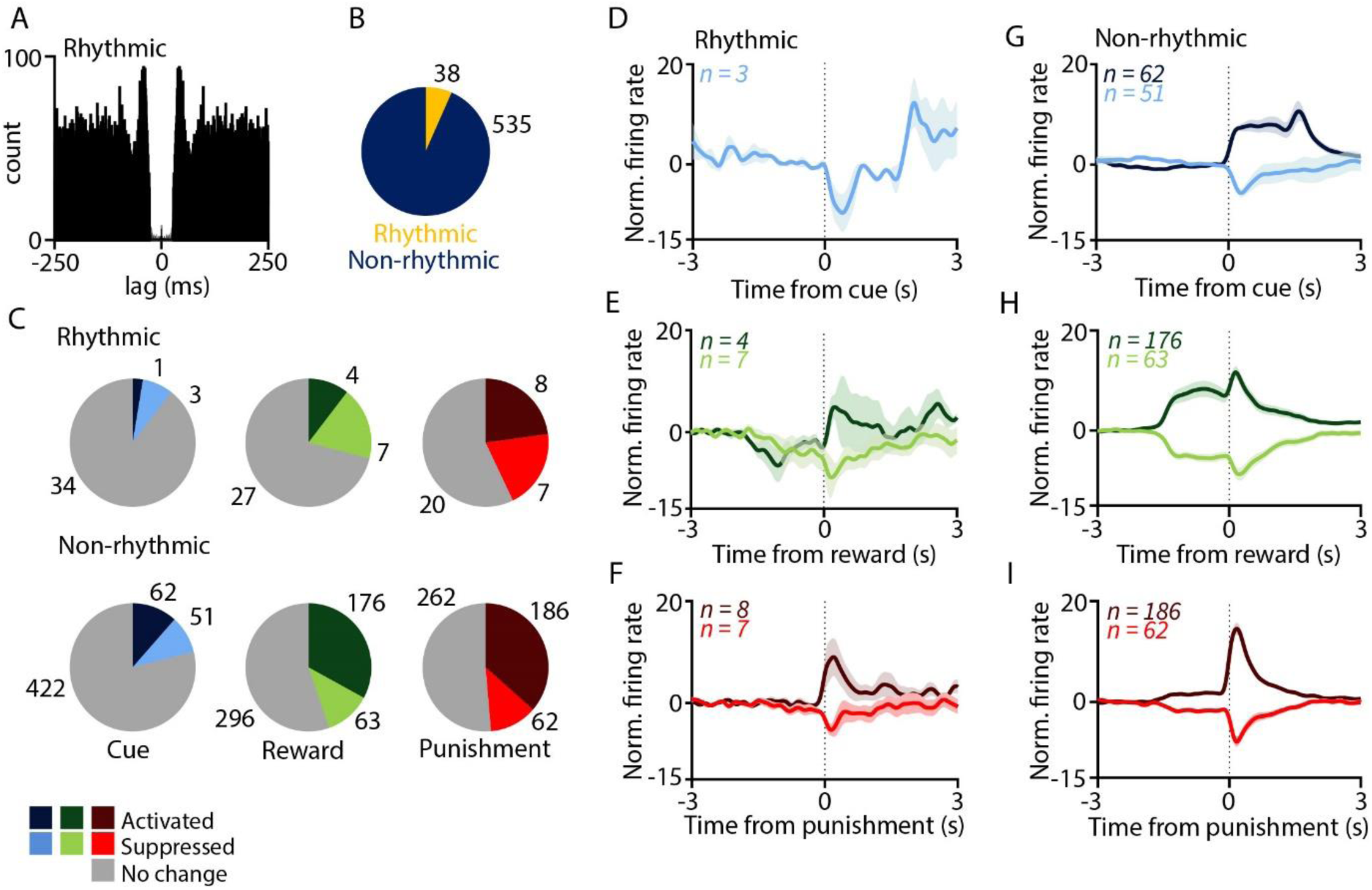
Salient events mostly recruit non-rhythmic neurons. ***A***, Example auto-correlogram of a rhythmically firing neuron. ***B***, Pie chart showing the proportion of rhythmic and non-rhythmic neurons. ***C***, Pie charts showing the number of rhythmic and non-rhythmic VP neurons activated and inhibited by cue, reward and punishment. ***D-I***, Average, z-scored PETHs of rhythmic ***(D-F)*** and non-non-rhythmic ***(G-I)*** VP neurons aligned to cue ***(D,G)*** reward ***(E,H)*** and punishment ***(F,I).***

### Indications of multiplexed burst and single spike code in the VP

Theoretical studies have suggested that since specific biophysical mechanisms are engaged to serve burst generation, burst firing may carry a representation independent from that of single spikes, creating a specific ‘burst code’ (Kepecs et al., 2002; Kepecs and Lisman, 2003). This may allow neurons to multiplex different sources of information; however, this idea has rarely been tested.

Therefore, we separated burst firing and single spike firing based on inter-spike interval (ISI) criteria. Specifically, bursts were defined by the first ISI <10ms and subsequent ISIs <15 ms (Royer et al., 2012; Laszlovszky et al., 2020). Next, bursts and single spikes of each neuron were aligned to behaviorally salient events. Burst and single spike firing often carried similar information about these events, indicated by correlated PETHs showing similar dynamics for bursts and single spikes (Fig. 6A-C). However, a subset of bursting VP neurons showed a dissociation of burst and single spike coding. The example neuron in Fig. 6D-F increased its firing rate after cue tone presentation. However, analysis of burst and single spike occurrence revealed that while single spike firing was elevated after the cues (Fig. 6F), burst firing showed a concurrent inhibition (Fig. 6E; this was not due to insufficient spike sorting). This suggests that a subset of bursting VP neurons exhibit separate representations of external events by bursts and single spikes, revealing a distinct ‘burst code’.

**Figure 6.**
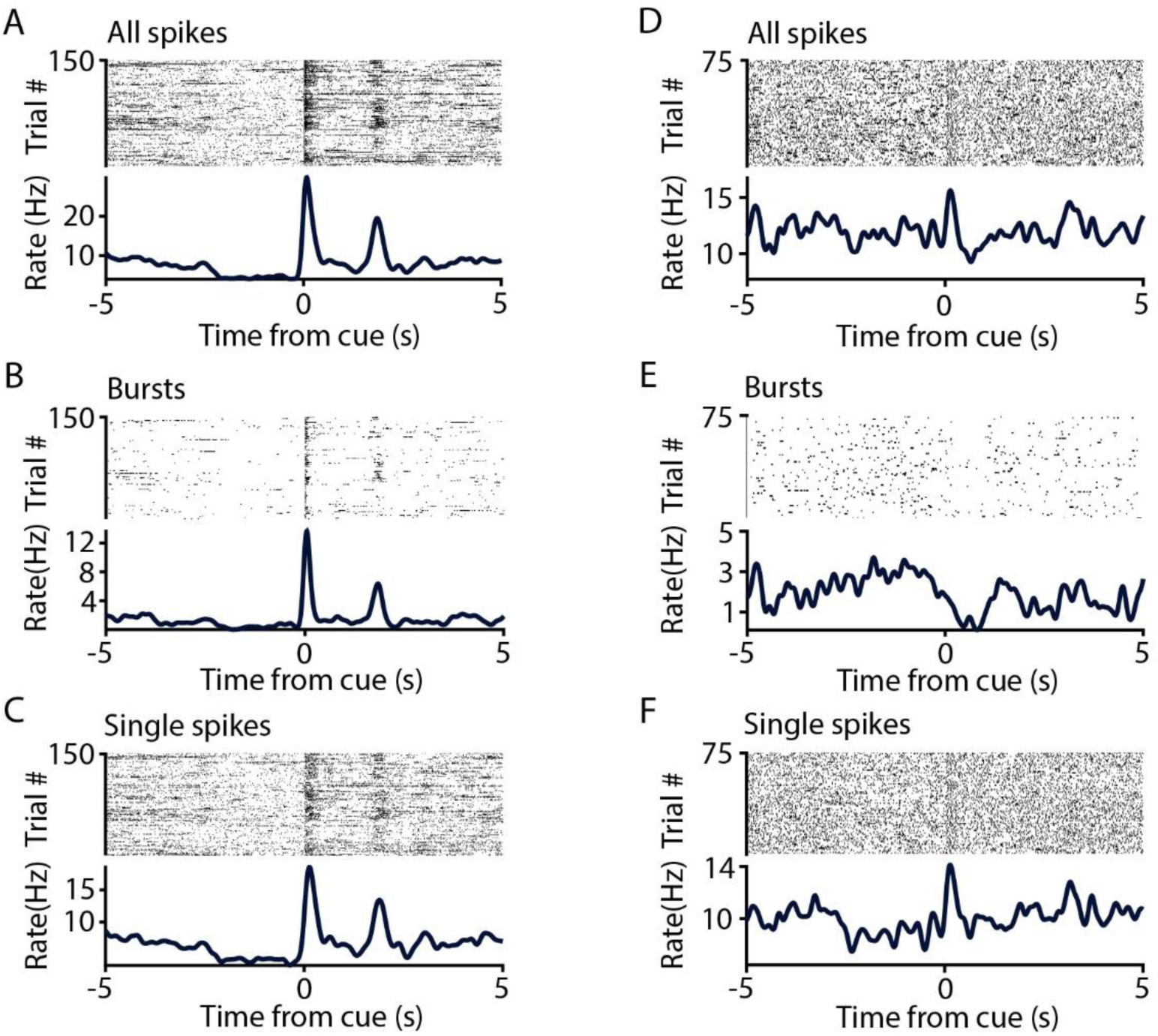
Dissociation of single spike and burst firing in a subset of VP neurons. ***A***, Example PETH of a VP neuron activated upon cue onset. The neuron shows increased burst ***(B)*** and single spike firing ***(C)*** after the cue. ***D***, Example of another cue-activated neuron, which shows decreased burst firing ***(E)*** and increased single spike firing ***(F)*** after cue presentation.

### Ventral pallidal neurons form synchronously firing assemblies

Neurons in some cortical and subcortical areas have been shown to form functional assemblies of co-firing cells (Fujisawa et al., 2008; Dupret et al., 2010). We performed a cross-correlation analysis of simultaneously recorded pairs of VP neurons (n = 795) and found many indications of functional connectivity. Neuronal pairs often showed a zero lag peak of cross-correlation typically taken as an indication of a common input. Narrow (1-2 ms wide) peaks within 1-4 ms from 0, on the other hand, usually indicate monosynaptic excitatory connections (Bartho et al., 2004; Fujisawa et al., 2008). We could identify small networks of VP neurons exhibiting pairwise synchrony, suggestive of assembly formation during associative learning in the ventral pallidum (Fig. 7.)

**Fig. 7.**
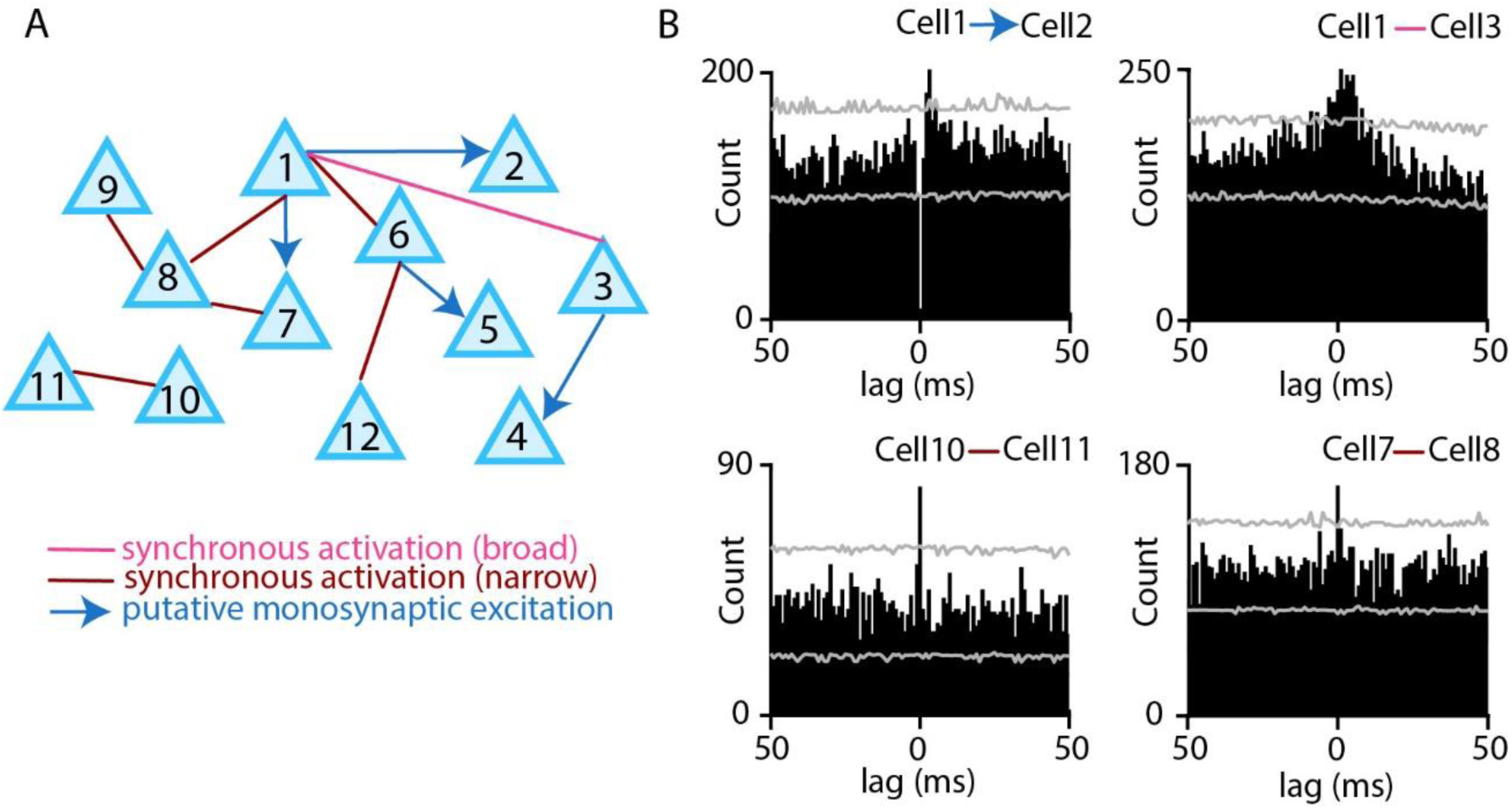
VP neurons form co-firing assemblies. ***A***, An example cell assembly formed by VP neurons during Pavlovian conditioning. Pink, narrow (1-2 ms) zero-lag synchrony; brown, broad (>=3 ms) zero-lag synchrony; blue, putative monosynaptic excitation. ***B***, Example cross-correlograms of neuronal pairs of this assembly. Grey, 95% confidence intervals.

### Neurons participating in assemblies respond more to reinforcement

Ventral pallidal neurons were sorted based on their cross-correlograms. Neurons that participated in synchronously firing assemblies based on significant zero-phase peaks detected in pairwise cross-correlations (Fig.7) were termed ‘synchronous neurons’, while neurons where no such concurrent activation was observed were called ‘asynchronous neurons’ (Fig.8A-B). While this distinction probably mis-labels some neurons that participate in assembly-formation as ‘asynchronous’ due to missed detections, we still uncovered prominent differences between the two groups. Neurons that participated in the detected assemblies (‘synchronous group’) showed more frequent responses to reward and punishment compared to those neurons for which we did not detect synchronous pairs (‘asynchronous group’; p = 0.0004, p = 0.009 for reward and punishment response, respectively; Chi square test). In contrast, the ‘asynchronous’ population showed a tendency to respond more to reward-predictive cues (Fig.8C-I; p = 0.02, chi square test). These differences probably represent an underestimation, since it is likely that we missed a fraction of synchronous activations.

**Fig. 8.**
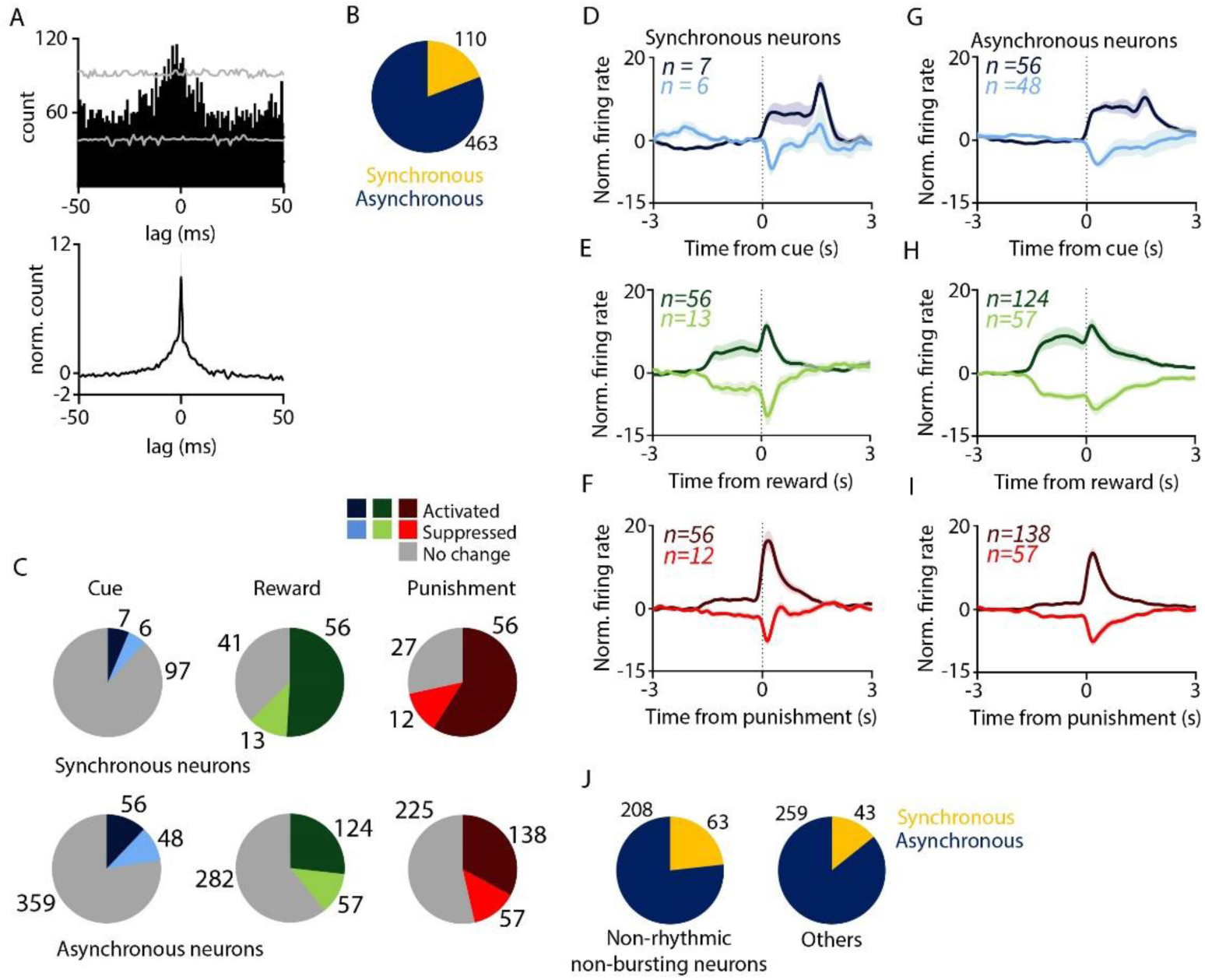
Synchronous neurons respond more to reinforcement. ***A***, Top, example cross-correlogram of synchronously activated neurons. Bottom, average cross-correlation of all synchronously active pairs. ***B***, Pie chart showing the proportion of synchronous and asynchronous VP neurons. ***C***, Pie charts showing the number of synchronous and asynchronous VP neurons activated and inhibited by cue, reward and punishment. ***D-I***, Average, z-scored PETHs of synchronous ***(D-F)*** and asynchronous ***(G-I)*** VP neurons aligned to cue ***(D,G)*** reward ***(E,H)*** and punishment ***(F,I). J***, Pie chart showing the proportion of synchronous neurons in the non-rhythmic-non-bursting population and in the rest of the population.

Since we found that non-bursting non-rhythmic neurons provide the majority of responses to reinforcement in the VP during Pavlovian learning, this finding also suggests that there may be more synchronously firing pairs of neurons in this non-bursting non-rhythmic population. We found that this indeed holds true (Fig.8J; p = 0.0056, Chi square test), suggesting the presence of a reinforcement responsive non-bursting-non-rhythmic VP population that forms co-active cell assemblies.

## Discussion

The role of VP in reward-related and motivated behavior has been extensively studied; however, there have been very few attempts to distinguish electrophysiologically defined neuronal populations during reinforcement learning (Avila and Lin, 2014; Kaplan et al., 2020). Therefore, the aim of this study was to characterize electrophysiologically distinct functional groups within the VP during reinforcement learning. We found that most responses to reward and punishment in the VP originated from a group of non-bursting-non-rhythmic neurons, suggesting that this population is dominant in representing reinforcers in the VP. Importantly, some bursting neurons showed differential responses when their bursts were contrasted with their single spikes, demonstrating that a specific ‘burst code’ may be present in the ventral pallidum (Kepecs and Lisman, 2003; Laszlovszky et al., 2020). VP neurons formed co-firing assemblies, and neurons participating in such assemblies were particularly responsive to reinforcement. In line with this result, we found that non-bursting non-rhythmic neurons were more likely to participate in co-firing assemblies. These results may suggest that an electrophysiologically defined population of the VP forms synchronous assemblies and transmit reinforcement signals to other areas.

Many studies have discussed the role of the VP in learning cue-reward associations (Tindell et al., 2004; Ito and Doya, 2009; Tachibana and Hikosaka, 2012; Avila and Lin, 2014; Richard et al., 2016a, 2018; Ahrens et al., 2018; Ottenheimer et al., 2018; Fujimoto et al., 2019) as well as in adapting behavioral responses to outcome during reinforcement learning (e.g. ‘liking’ reactions after reward delivery and ‘disgust’ reactions to aversive stimuli) (Smith and Berridge, 2005; Tindell et al., 2006; Ho and Berridge, 2014), gated by internal state (Chang et al., 2017; Stephenson-Jones et al., 2020). Many of these experiments only included rewarded and omitted trials, whereas comparatively fewer papers featured aversive stimuli (Knowland et al., 2017; Saga et al., 2017; Wulff et al., 2019; Kaplan et al., 2020; Stephenson-Jones et al., 2020). Cued reward size modifications were often included (Tachibana and Hikosaka, 2012; Stephenson-Jones et al., 2020), although how the VP adapts to probabilistic cues that are notoriously harder to learn, since they require integration over many trials, has remained largely unexplored.

Here we trained mice on a Pavlovian reinforcement learning task where we incorporated both reward and punishment into our task design, in order to examine VP neuronal activity patterns upon both positive and negative outcomes. We found a VP population activated by both reward and reward-predicting cues, consistent with previous findings (Tindell et al., 2004; Ahrens et al., 2016, 2018; Stephenson-Jones et al., 2020). However, we found even more punishment-responsive VP neurons that have largely been overlooked before. Moreover, responses to punishment were significantly faster than reward or cue-elicited firing rate changes.

A significant population showed inhibition to cues and reinforcement regardless of valence, consistent with Type III. GABAergic neurons in the study by Stephenson and colleagues (Stephenson-Jones et al., 2020). However, we found very few neurons that responded with opposite firing rate changes to reward and punishment, unlike in the above cited report, and in most cases the signs of responses were correlated across stimuli (Fig.2O). An important factor that likely underlies these differences is that we used a probabilistic task design, in which both cues were followed by reward, punishment or nothing with different, set probabilities. Thus, probabilistic expectations provide a task context in which VP neurons tend to respond more positively, as all cues carry some positive value, also indicated by the dominance of neuronal activation vs. inhibition in our recordings. Additionally, a number of these cells might be modulated by incentive salience rather than encoding outcome valence, as found in previous studies (Tindell et al., 2004, 2009; Ahrens et al., 2018; Stephenson-Jones et al., 2020). We also note that most of our recordings originated from the anterior half of the VP, thus known anatomical differences along the antero-posterior axis (Stratford et al., 1999; Mahler et al., 2014) may have contributed to some of these differences.

The VP is a key node in the integration of limbic and motor processes (Fujimoto et al., 2019). The probabilistic outcome contingencies of our Pavlovian task enabled us to show that VP neurons’ cue responses are modulated by reward expectation. To a lesser extent, outcome responses were also modulated based on being expected or surprising. Reward prediction error responses are expected to occur after reward-predictive sensory cues and diminish for anticipated vs. unexpected rewards (Schultz et al., 1997; Kim et al., 2019). In contrast, we found slightly but significantly higher firing rate elevation after expected than surprising rewards. This argues against a prominent reward prediction error coding in probabilistic cued reward scenarios in the VP. At the same time, this finding may be consistent with representing incentive salience, inasmuch as reward anticipation may allow the limbic-motor system to better anticipate the time of reward and boost its response by preparatory activities. A recent study elegantly showed that different subpopulations of VP neurons may be responsible for representing incentive salience and reward prediction errors, adding and another layer of complexity to information multiplexing in the VP (Kaplan et al., 2020).

The VP is known to be strongly innervated by dopaminergic fibers arising from the ventral tegmental area (VTA). It appears that the functional role of this connection is the modulation of locomotion by acting on VP neuronal output (Klitenick et al., 1992). Morevoer, the VTA itself is also considered to be crucial for reward expectation coding (Hollerman and Schultz, 1998) and has been shown to promote place preference via its afferent projections from the VP (Faget et al., 2018). This strong reciprocal connection between the two areas could serve as a neural basis of reward seeking behavior in rodents. We should note however, that the VP also lies at the intersection of basal ganglia and basal forebrain circuits, the latter also featuring prominent reward-prediction activity and salience coding (Lin and Nicolelis, 2008; Avila and Lin, 2014; Hangya et al., 2015); therefore, multiple origins of such signals are feasible. In this regard, an important finding showed that VP responses to reward appear earlier than those in the nucleus accumbens, making the previously hypothesized accumbens to VP information transfer less likely (Richard et al., 2016b; Ottenheimer et al., 2018).

We characterized the activity of bursting VP neurons and found that they were less responsive to reward and punishment than non-bursting neurons. This was surprising, as bursts of action potentials are often thought to be associated with stronger excitatory drive that may lead to larger firing rate increases. Indeed, in the basal forebrain, bursting neurons as well as burst responses were associated with populations responsive to salient stimuli (Lin and Nicolelis, 2008; Laszlovszky et al., 2020). In addition, our findings are seemingly at odds with Avila and Lin demonstrating burst responses to salient events in the VP (Avila and Lin, 2014). However, we based our definition on classical burst criteria in earlier studies (Pang et al., 1998; Royer et al., 2012), while Avila and Lin chose an elegant unbiased clustering method, after which a group of bursting neurons emerged automatically. Bursting in those neurons was, however, substantially slower (15-30 ms); therefore, these two ‘bursting’ populations may in fact be non-overlapping. As this confusion of terminology is rampant in the subcortical literature, we suggest introducing the terms ‘fast-burst’ and ‘slow-burst’ to differentiate between interspike-intervals shorter or longer than 10 ms.

Different higher order firing patterns may represent specific information, as a special case of temporal code (Panzeri et al., 2010). Thus, bursts of action potentials conceivably code different variables from single spikes even within single neurons, allowing within-cell multiplexing of information (Kepecs et al., 2002; Kepecs and Lisman, 2003). For instance, bursts of visual thalamic neurons were shown to have sharper tuning than single spikes (Reinagel et al., 1999), and basal forebrain bursts of both cholinergic and non-cholinergic neurons represent specific information about salient stimuli (Lin and Nicolelis, 2008; Hangya et al., 2015; Laszlovszky et al., 2020). Even ‘slow-bursts’ and ‘fast-bursts’ may exhibit different properties, possibly reconciling different results mentioned above (Avila and Lin, 2014). In accordance, we found VP neurons that showed strong differences in their burst and single spike occurrence after salient stimuli, which could in some cases change in opposite directions (Fig. 6.)

Pang and colleagues identified a fast rhythmic type of VP neuron, with rhythmicity frequency in the beta/gamma bands (Fig. 8A in (Pang et al., 1998)). Recently, VP gamma activity was linked to somatostatin (SOM)-expressing GABAergic neurons that influenced movement speed (Espinosa et al., 2019). In contrast, SOM neurons in the medial septum did not exhibit gamma-correlation. Indeed, in other parts of the basal forebrain, gamma oscillations were better correlated with parvalbumin-expressing GABAergric neurons (Kim et al., 2015). This basal forebrain-VP dissociation posits that these SOM GABAergic VP neurons may be more linked to basal ganglia than basal forebrain activity. Consistent with this, we found that these fast rhythmic neurons are not prominent contributors of VP reinforcement responses.

The presence of cell assemblies have been previously demonstrated in the hippocampus, nucleus accumbens and basal forebrain (Harris et al., 2003). Dynamically forming cell assemblies of the hippocampus (Harris et al., 2003; Tingley et al., 2015; Trouche et al., 2019) were linked to spatial navigation and episodic memory recall (Pastalkova et al., 2008; Dupret et al., 2010; Mamad et al., 2017). Assemblies in the basal forebrain were suggested to organize behavior in an attention task (Tingley et al., 2014, 2015). We demonstrated that synchronously firing cell assemblies are also formed in the VP during Pavlovian conditioning. Moreover, the temporal scale of co-firing closely matched the 10-30 ms previously suggested to be a conserved parameter under biophysical constraints (Harris et al., 2003) (Fig.8A). Consistent with the idea that ‘transient synchrony of anatomically distributed groups of neurons underlies processing of both external sensory input and internal cognitive mechanisms’ (Harris et al., 2003), we found that neurons participating in co-firing ensembles are more responsive to reward and punishment during Pavlovian learning.

The output of VP neurons can be modulated by enkephalins and dopamine, since VP neurons express µ and κ opioid receptors as well as D1 and D2 receptors (Panagis et al., 1998; Kupchik et al., 2015; Clark and Bracci, 2018). This receptor pattern makes the VP especially sensitive to drugs targeting the dopaminergic and opioid system, including opiates and cocaine (Mickiewicz et al., 2009; Mahler et al., 2014; Creed et al., 2016; Heinsbroek et al., 2020). A shift in the activity pattern of VP neurons can lead to maladaptive behavior, such as perseveration or intracranial self-stimulation in rodents, or severe addiction in humans (Hubner and Koob, 1990; Ottenheimer et al., 2019). Moreover, in patients with long term addiction, the pathological function is accompanied by morphological changes (Müller et al., 2019). Therefore, the VP might be an ideal target for the treatment of drug addiction and habitual relapse. The nucleus accumbens, one of the main inputs to the VP, has already been proposed as a potential target for deep brain stimulation (DBS) in patients with addiction (Kuhn et al., 2014; Müller et al., 2015). However, the central position of the VP in the reward circuitry and its strong relation to addiction could make the VP another plausible target for DBS (Yu et al., 2016; Mahoney et al., 2018). Gaining a better foothold on understanding the activity patterns and coding schemes of the VP will be fundamental for developing an effective stimulation protocol while minimizing side effects.

## Acknowledgement

This work was supported by the ‘Lendület’ Program of the Hungarian Academy of Sciences (LP2015-2/2015), NKFIH KH125294 and the European Research Council Starting Grant no. 715043. We thank Katalin Lengyel for her help with histology and Katalin Sviatkó and Sergio Martínez-Bellver for their help with behavioral training.

## Author contributions

BH developed the idea and conceptualized the manuscript. PH and JH performed the experiments. PH, BH and JH performed data analysis. PH generated the figures. BH and PH wrote the manuscript with input from JH.

## References

Ahrens AM, Ferguson LM, Robinson TE, Aldridge JW (2018) Dynamic encoding of incentive salience in the ventral pallidum: Dependence on the form of the reward cue. eNeuro 5:1–16.

Ahrens AM, Meyer PJ, Ferguson LM, Robinson TE, Wayne Aldridge J (2016) Neural activity in the ventral pallidum encodes variation in the incentive value of a reward cue. J Neurosci 36:7957–7970.

Avila I, Lin S-C (2014) Distinct neuronal populations in the basal forebrain encode motivational salience and movement. Front Behav Neurosci 8:421.

Bartho P, Hirase H, Monconduit L, Zugaro M, Harris KD, Buzsaki G (2004) Characterization of neocortical principal cells and interneurons by network interactions and extracellular features. J Neurophysiol 92:600–608.

Chang SE, Smedley EB, Stansfield KJ, Stott JJ, Smith KS (2017) Optogenetic inhibition of ventral pallidum neurons impairs context-driven salt seeking. J Neurosci 37:5670–5680.

Clark M, Bracci E (2018) Dichotomous Dopaminergic Control of Ventral Pallidum Neurons. 12:1–19.

Creed M, Ntamati NR, Chandra R, Lobo MK, Lüscher C (2016) Convergence of Reinforcing and Anhedonic Cocaine Effects in the Ventral Pallidum. Neuron 92:214–226.

Dupret D, O’Neill J, Pleydell-Bouverie B, Csicsvari J (2010) The reorganization and reactivation of hippocampal maps predict spatial memory performance. Nat Neurosci 13:995–1002.

Espinosa N, Alonso A, Lara-Vasquez A, Fuentealba P (2019) Basal forebrain somatostatin cells differentially regulate local gamma oscillations and functionally segregate motor and cognitive circuits. Sci Rep 9:1–12.

Faget L, Zell V, Souter E, McPherson A, Ressler R, Gutierrez-Reed N, Yoo JH, Dulcis D, Hnasko TS (2018) Opponent control of behavioral reinforcement by inhibitory and excitatory projections from the ventral pallidum. Nat Commun 9:849.

Fuhrmann F, Justus D, Sosulina L, Kaneko H, Beutel T, Friedrichs D, Schoch S, Schwarz MK, Fuhrmann M, Remy S (2015) Locomotion, Theta Oscillations, and the Speed-Correlated Firing of Hippocampal Neurons Are Controlled by a Medial Septal Glutamatergic Circuit. Neuron 86:1253–1264.

Fujimoto A, Hori Y, Nagai Y, Kikuchi E, Oyama K, Suhara T, Minamimoto T (2019) Signaling incentive and drive in the primate ventral pallidum for motivational control of goal-directed action. J Neurosci 39:1793–1804.

Fujisawa S, Amarasingham A, Harrison MT, Buzsáki G (2008) Behavior-dependent short-term assembly dynamics in the medial prefrontal cortex. Nat Neurosci 11:823–833.

Hangya B, Li Y, Muller RU, Czurkó A (2010) Complementary spatial firing in place cell-interneuron pairs. J Physiol 588:4165–4175.

Hangya B, Ranade SP, Lorenc M, Kepecs A (2015) Central Cholinergic Neurons Are Rapidly Recruited by Reinforcement Feedback. Cell 162:1155–1168.

Harris KD, Csicsvari J, Hirase H, Dragoi G, Buzsáki G (2003) Organization of cell assemblies in the hippocampus. Nature 424:552–556.

Heinsbroek JA, Bobadilla AC, Dereschewitz E, Assali A, Chalhoub RM, Cowan CW, Kalivas PW (2020) Opposing Regulation of Cocaine Seeking by Glutamate and GABA Neurons in the Ventral Pallidum. Cell Rep 30:2018-2027.e3.

Ho C, Berridge KC (2014) Excessive disgust caused by brain lesions or temporary inactivations : mapping hotspots of the nucleus accumbens and ventral pallidum. 40:3556–3572.

Hollerman JR, Schultz W (1998) Dopamine neurons report an error in the temporal prediction of reward during learning. Nat Neurosci 1:304–309.

Hubner CB, Koob GF (1990) The ventral pallidum plays a role in mediating cocaine and heroin self-administration in the rat. Brain Res 508:20–29.

Ito M, Doya K (2009) Validation of decision-making models and analysis of decision variables in the rat basal ganglia. J Neurosci 29:9861–9874.

Kaplan A, Mizrahi-Kliger AD, Israel Z, Adler A, Bergman H (2020) Dissociable roles of ventral pallidum neurons in the basal ganglia reinforcement learning network. Nat Neurosci 23:556–564.

Kepecs A, Lisman J (2003) Information encoding and computation with spikes and bursts. Netw Comput Neural Syst 14:103–118.

Kepecs A, Wang X-J, Lisman J (2002) Bursting neurons signal input slope. J Neurosci 22:9053–9062.

Kim HR, Malik AN, Mikhael JG, Bech P, Tsutsui-Kimura I, Sun F, Zhang Y, Li Y, Watabe-Uchida M, Gershman SJ, Uchida N (2019) A unified framework for dopamine signals across timescales. bioRxiv:803437.

Kim T, Thankachan S, Mckenna JT, Mcnally JM, Yang C, Hyun J (2015) Cortically projecting basal forebrain parvalbumin neurons regulate cortical gamma band oscillations. 112:3535–3540.

Klitenick MA, Deutch AY, Churchill L, Kalivas PW (1992) Topography and functional role of dopaminergic projections from the ventral mesencephalic tegmentum to the ventral pallidum. Neuroscience 50:371–386.

Knowland D, Lilascharoen V, Pacia CP, Shin S, Wang EHJ, Lim BK (2017) Distinct Ventral Pallidal Neural Populations Mediate Separate Symptoms of Depression. Cell 170:284-297.e18.

Kuhn J, Möller M, Treppmann JF, Bartsch C, Lenartz D, Gruendler TOJ, Maarouf M, Brosig A, Barnikol UB, Klosterkötter J, Sturm V (2014) Deep brain stimulation of the nucleus accumbens and its usefulness in severe opioid addiction. Mol Psychiatry 19:145–146.

Kupchik YM, Brown RM, Heinsbroek JA, Lobo MK, Schwartz DJ, Kalivas PW (2015) Coding the direct/indirect pathways by D1 and D2 receptors is not valid for accumbens projections. Nat Neurosci 18:1230–1232.

Kvitsiani D, Ranade S, Hangya B, Taniguchi H, Huang JZ, Kepecs A (2013) Distinct behavioural and network correlates of two interneuron types in prefrontal cortex. Nature 498:363–366.

Laszlovszky T, Schlingloff D, Hegedüs P, Freund TF, Gulyás A, Kepecs A, Hangya B (2020) Distinct synchronization, cortical coupling and behavioural function of two basal forebrain cholinergic neuron types. bioRxiv:703090.

Lin S-C, Nicolelis M a L (2008) Neuronal ensemble bursting in the basal forebrain encodes salience irrespective of valence. Neuron 59:138–149.

Mahler S V., Vazey EM, Beckley JT, Keistler CR, Mcglinchey EM, Kaufling J, Wilson SP, Deisseroth K, Woodward JJ, Aston-Jones G (2014) Designer receptors show role for ventral pallidum input to ventral tegmental area in cocaine seeking. Nat Neurosci 17:577–585.

Mahoney EC, Zeng A, Yu W, Rowe M, Sahai S, Feustel PJ, Ramirez-Zamora A, Pilitsis JG, Shin DS (2018) Ventral pallidum deep brain stimulation attenuates acute partial, generalized and tonic-clonic seizures in two rat models. Epilepsy Res 142:36–44.

Mamad O, Stumpp L, McNamara HM, Ramakrishnan C, Deisseroth K, Reilly RB, Tsanov M (2017) Place field assembly distribution encodes preferred locations Csicsvari J, ed. PLOS Biol 15:e2002365.

Maurice N, Deniau JM, Menetrey A, Glowinski J, Thierry AM (1997) Position of the ventral pallidum in the rat prefrontal cortex-basal ganglia circuit. Neuroscience 80:523–534.

Mickiewicz AL, Dallimore JE, Napier TC (2009) The ventral pallidum is critically involved in the development and expression of morphine-induced sensitization. Neuropsychopharmacology 34:874–886.

Müller UJ, Mawrin C, Frodl T, Dobrowolny H, Busse S, Bernstein H-G, Bogerts B, Truebner K, Steiner J (2019) Reduced volumes of the external and internal globus pallidus in male heroin addicts: a postmortem study. Eur Arch Psychiatry Clin Neurosci 269:317–324.

Müller UJ, Truebner K, Schiltz K, Kuhn J, Mawrin C, Dobrowolny H, Bernstein HG, Bogerts B, Steiner J (2015) Postmortem volumetric analysis of the nucleus accumbens in male heroin addicts : implications for deep brain stimulation. Eur Arch Psychiatry Clin Neurosci 265:647–653.

Ottenheimer D, Richard JM, Janak PH (2018) Ventral pallidum encodes relative reward value earlier and more robustly than nucleus accumbens. Nat Commun 9:4350.

Ottenheimer DJ, Wang K, Haimbaugh A, Janak PH, Richard JM (2019) Recruitment and disruption of ventral pallidal cue encoding during alcohol seeking. Eur J Neurosci 50:3428–3444.

Panagis G, Kastellakis A, Spyraki C (1998) Involvement of the ventral tegmental area opiate receptors in self-stimulation elicited from the ventral pallidum. Psychopharmacology (Berl) 139:222–229.

Pang K, Tepper JM, Zaborszky L (1998) Morphological and Electrophysiological Characteristics of Noncholinergic Basal. 204:186–204.

Panzeri S, Brunel N, Logothetis NK, Kayser C (2010) Sensory neural codes using multiplexed temporal scales. Trends Neurosci 33:111–120.

Pastalkova E, Itskov V, Amarasingham A, Buzsáki G (2008) Internally generated cell assembly sequences in the rat hippocampus. Science 321:1322–1327.

Paxinos G, Franklin KBJ, Franklin KBJ (2001) The mouse brain in stereotaxic coordinates. Academic Press.

Prasad AA, Xie C, Chaichim C, Nguyen JH, McClusky HE, Killcross S, Power JM, McNally GP (2020) Complementary roles for ventral pallidum cell types and their projections in relapse. J Neurosci 40:880–893.

Reinagel P, Godwin D, Sherman SM, Koch C (1999) Encoding of Visual Information by LGN Bursts. J Neurophysiol 81:2558–2569.

Richard JM, Ambroggi F, Janak PH, Fields HL (2016a) Ventral Pallidum Neurons Encode Incentive Value and Promote Cue-Elicited Instrumental Actions. Neuron 90:1165–1173.

Richard JM, Ambroggi F, Janak PH, Fields HL (2016b) Ventral Pallidum Neurons Encode Incentive Value and Promote Cue-Elicited Instrumental Actions. Neuron 90:1165–1173.

Richard JM, Stout N, Acs D, Janak PH (2018) Ventral pallidal encoding of reward-seeking behavior depends on the underlying associative structure. Elife 7:1–25.

Root DH, Melendez RI, Zaborszky L, Napier TC (2015) The ventral pallidum: Subregion-specific functional anatomy and roles in motivated behaviors. Prog Neurobiol 130:29–70.

Royer S, Zemelman B V, Losonczy A, Kim J, Chance F, Magee JC, Buzsáki G (2012) Control of timing, rate and bursts of hippocampal place cells by dendritic and somatic inhibition. Nat Neurosci 15:769–775.

Saga Y, Richard A, Sgambato-Faure V, Hoshi E, Tobler PN, Tremblay L (2017) Ventral pallidum encodes contextual information and controls aversive behaviors. Cereb Cortex 27:2528–2543.

Schmitzer-Torbert N, Jackson J, Henze D, Harris K, Redish a D (2005) Quantitative measures of cluster quality for use in extracellular recordings. Neuroscience 131:1–11.

Schultz W, Dayan P, Montague PR (1997) A neural substrate of prediction and reward. Science 275:1593–1599.

Smith KS, Berridge KC (2005) The ventral pallidum and hedonic reward: Neurochemical maps of sucrose “liking” and food intake. J Neurosci 25:8637–8649.

Smith KS, Tindell AJ, Aldridge JW, Berridge KC (2009) Ventral pallidum roles in reward and motivation. Behav Brain Res 196:155–167.

Solari N, Sviatkó K, Laszlovszky T, Hegedüs P, Hangya B (2018) Open Source Tools for Temporally Controlled Rodent Behavior Suitable for Electrophysiology and Optogenetic Manipulations. Front Syst Neurosci 12:18.

Stephenson-Jones M, Bravo-Rivera C, Ahrens S, Furlan A, Xiao X, Fernandes-Henriques C, Li B (2020) Opposing Contributions of GABAergic and Glutamatergic Ventral Pallidal Neurons to Motivational Behaviors. Neuron 105:921-933.e5.

Stratford TR, Kelley AE, Simansky KJ (1999) Blockade of GABA(A) receptors in the medial ventral pallidum elicits feeding in satiated rats. Brain Res 825:199–203.

Tachibana Y, Hikosaka O (2012) The Primate Ventral Pallidum Encodes Expected Reward Value and Regulates Motor Action. Neuron 76:826–837.

Tindell AJ, Berridge KC, Aldridge JW (2004) Ventral Pallidal Representation of Pavlovian Cues and Reward : Population and Rate Codes. 24:1058–1069.

Tindell AJ, Smith KS, Berridge KC, Aldridge JW (2009) Dynamic computation of incentive salience: “Wanting” what was never “liked.” J Neurosci 29:12220–12228.

Tindell AJ, Smith KS, Peciña S, Berridge KC, Aldridge JW (2006) Ventral pallidum firing codes hedonic reward: when a bad taste turns good. J Neurophysiol 96:2399–2409.

Tingley D, Alexander AS, Kolbu S, de Sa VR, Chiba AA, Nitz DA (2014) Task-phase-specific dynamics of basal forebrain neuronal ensembles. Front Syst Neurosci 8:1–15.

Tingley D, Alexander AS, Quinn LK, Chiba AA, Nitz DA (2015) Cell Assemblies of the Basal Forebrain. J Neurosci 35:2992–3000.

Trouche S, Koren V, Doig NM, Ellender TJ, El-Gaby M, Lopes-dos-Santos V, Reeve HM, Perestenko P V., Garas FN, Magill PJ, Sharott A, Dupret D (2019) A Hippocampus-Accumbens Tripartite Neuronal Motif Guides Appetitive Memory in Space. Cell 176:1393-1406.e16.

van den Bos R, Cools AR (1991) Motor activity and the GABAA-receptor in the ventral pallidum/substantia innominata complex. Neurosci Lett 124:246–250.

Wulff AB, Tooley J, Marconi LJ, Creed MC (2019) Ventral pallidal modulation of aversion processing. Brain Res 1713:62–69.

Yang Y, Cui Y, Sang K, Dong Y, Ni Z, Ma S, Hu H (2018) Ketamine blocks bursting in the lateral habenula to rapidly relieve depression. Nature 554:317–322.

Yu W, Walling I, Smith AB, Ramirez-Zamora A, Pilitsis JG, Shin DS (2016) Deep Brain Stimulation of the Ventral Pallidum Attenuates Epileptiform Activity and Seizing Behavior in Pilocarpine-Treated Rats. Brain Stimul 9:285–295.

Záborszky L, Cullinan WE (1992) Projections from the nucleus accumbens to cholinergic neurons of the ventral pallidum: a correlated light and electron microscopic double-immunolabeling study in rat. Brain Res 570:92–101.

Zaborszky L, Pol A Van Den, Gyengesi E (2012) The Basal Forebrain Cholinergic Projection System in Mice. In: The mouse nervous system, 1st ed. (Watson C, Paxinos G, Puelles L, eds), pp 684–718. Amsterdam: Elsevier.

Zahm DS, Williams E, Wohltmann C (1996) Ventral striatopallidothalamic projection: IV. Relative involvements of neurochemically distinct subterritories in the ventral pallidum and adjacent parts of the rostroventral forebrain. J Comp Neurol 364:340–362.

